# Isolation of Genomic Deoxyxylulose Phosphate Reductoisomerase (DXR) Mutations Conferring Resistance to Fosmidomycin

**DOI:** 10.1101/296954

**Authors:** Gur Pines, Marcelo C. Bassalo, Eun Joong Oh, Alaksh Choudhury, Andrew D. Garst, Ryan T. Gill

## Abstract

Sequence to activity mapping technologies are rapidly developing, enabling the isolation of mutations that confer novel phenotypes. Here we used the CRISPR EnAbled Trackable genome Engineering (CREATE) technology to investigate the inhibition of the essential *IspC* gene in *Escherichia coli*. *IspC* gene product, Deoxyxylulose Phosphate Reductoisomerase (DXR), converts 1-deoxy-D-xylulose 5-phosphate to 2-C-methyl-D-erythritol 4-phosphate in the DXP pathway. Since this pathway is shared with many pathogenic bacteria and protozoa and is missing in humans, it is an appealing target for inhibition. We created a full saturation library of 33 sites proximal to ligand binding and other sites and challenged it with the DXR-specific inhibitor, fosmidomycin. We identified several mutations that confer fosmidomycin resistance. All sites are highly conserved and also exist in pathogens including the malaria-inducing *Plasmodium falciparum*. These findings may have general implications on the isolation of resistance-conferring mutations and specifically, may affect the design of future generations of fosmidomycin-based drugs.

**Significance:** The emergence of acquired drug resistance is a natural process that is likely to occur under most circumstances. Recently-developed technologies allow to map relative fitness contribution of multiple mutations in parallel. Such approaches may be used to predict which mutations are most likely to confer resistance, instead of waiting for them to evolve spontaneously. In this study, a rationally-designed *IspC* mutant library was generated genomically in *E. coli*. Mutants resistant to fosmidomycin, an antimalarial drug were identified, and most were in the highly conserved proline at position 274. These results may have implications on next-generation fosmidomycin drug design, and more broadly, this approach may be used for predicting mutational acquired resistance.

## Introduction

Drug resistance is currently a major concern in relation to human health. It emerges when using antibiotics to battle a bacterial infection, while treating cancer, or when using pesticides or herbicides. Resistance may arise through multiple mechanisms including the horizontal transfer of resistance genes, differential expression of various genes such as efflux pumps, and through mutations of the target or other genes (1, 2). Mutational resistance is an evolutionary process that takes advantage of increased mutational rates triggered by genome instability (3, 4) or stress signals (5–7), scanning the mutational landscape for an adaptive solution. The reduced pace of new antibiotic discovery, combined with widespread use of current antibiotics led to numerous fatalities due to multidrug-resistant bacterial infections suggesting that we are entering a post-antibiotic age (8–10). Similarly, tumor cells frequently mutate the gene target of personalized medicine and thus achieve resistance (11–13). Hence, the ability to predict which mutations will confer resistance to a particular drug is a critical early step in the drug development pipeline.

Sequence to activity mapping approaches explore the mutational landscape by assessing the impact of mutations on fitness. Such approaches link methods for producing sequence diversity with selection or screening techniques to isolate phenotypes of interest. Sequence diversity may be generated randomly, for example by error-prone PCR or adaptive laboratory evolution experiments. However, these approaches are not systematic and might miss important information. Saturation mutagenesis experiments are more comprehensive but require some a priori knowledge about the exact sites to be mutated. Recent advances in large-scale DNA synthesis and sequencing allow more thorough investigations by saturating every site within the target gene (14–18).

Deoxyxylulose Phosphate Reductoisomerase (DXR) catalyzes the first committed step of DXP metabolic pathway and converts 1-deoxy-D-xylulose 5-phosphate (DOXP) to 2-C-methyl-D-erythritol 4-phosphate (MEP). The DXP pathway produces isopentenyl pyrophosphate (IPP) and dimethylallyl pyrophosphate (DMAPP), which are essential to all life forms and the precursors of the broadly important isoprenoid pathway. While most eukaryotes use the mevalonate pathway to produce these molecules, bacteria, and some protozoa use the alternative DXP pathway (19). Thus, the DXP pathway is an attractive target for inhibition (20, 21). Indeed, the DXR inhibitor fosmidomycin (FSM) was successfully used to treat malaria patients either as a monotherapy or in combination with other drugs (22–24). Several pathways were found for FSM resistance in various organisms, including lack of uptake and DXR amplification (25–27). To date, a single point mutation in the *E. coli* DXR gene was found to confer FSM resistance (28).

We recently reported on a genome editing method that combines recombineering and CRISPR-based methods to produce large-scale libraries with a single amino acid polymorphism resolution (14). In short, the CRISPR EnAbled Trackable genome Engineering (CREATE) approach uses plasmid-based homology arms containing the desired mutation and another silent PAM mutation that protects the edited genome from further digestion (Fig. 1A). Synthesis of multiple such homology arms allows the pooled cloning of plasmid libraries that are then introduced to the target cells (Fig. 1B-D). Unedited wild-type genomes are attacked by the CRISPR-CAS9 system, inducing genomic double-strand breaks and subsequent cell death, while edited cells are protected thanks to the PAM mutation. Relative fitness values can be calculated by deep sequencing of the editing plasmids that serve as barcodes for mutation identification. Here, we map the sequence to activity relationships of 33 sites in the *E. coli* DXR gene, *ispC*. We challenged the edited cells with FSM and isolated several previously uncharacterized mutations that confer resistance. We also mapped the probability of each resistance-conferring mutation to occur according to the required number of base substitutions. These results may prove valuable when designing future drugs that are based on FSM structure. Moreover, the approach described here is broadly applicable to map many other resistance phenotypes with potential implications on drug development timelines.

**Fig. 1.**
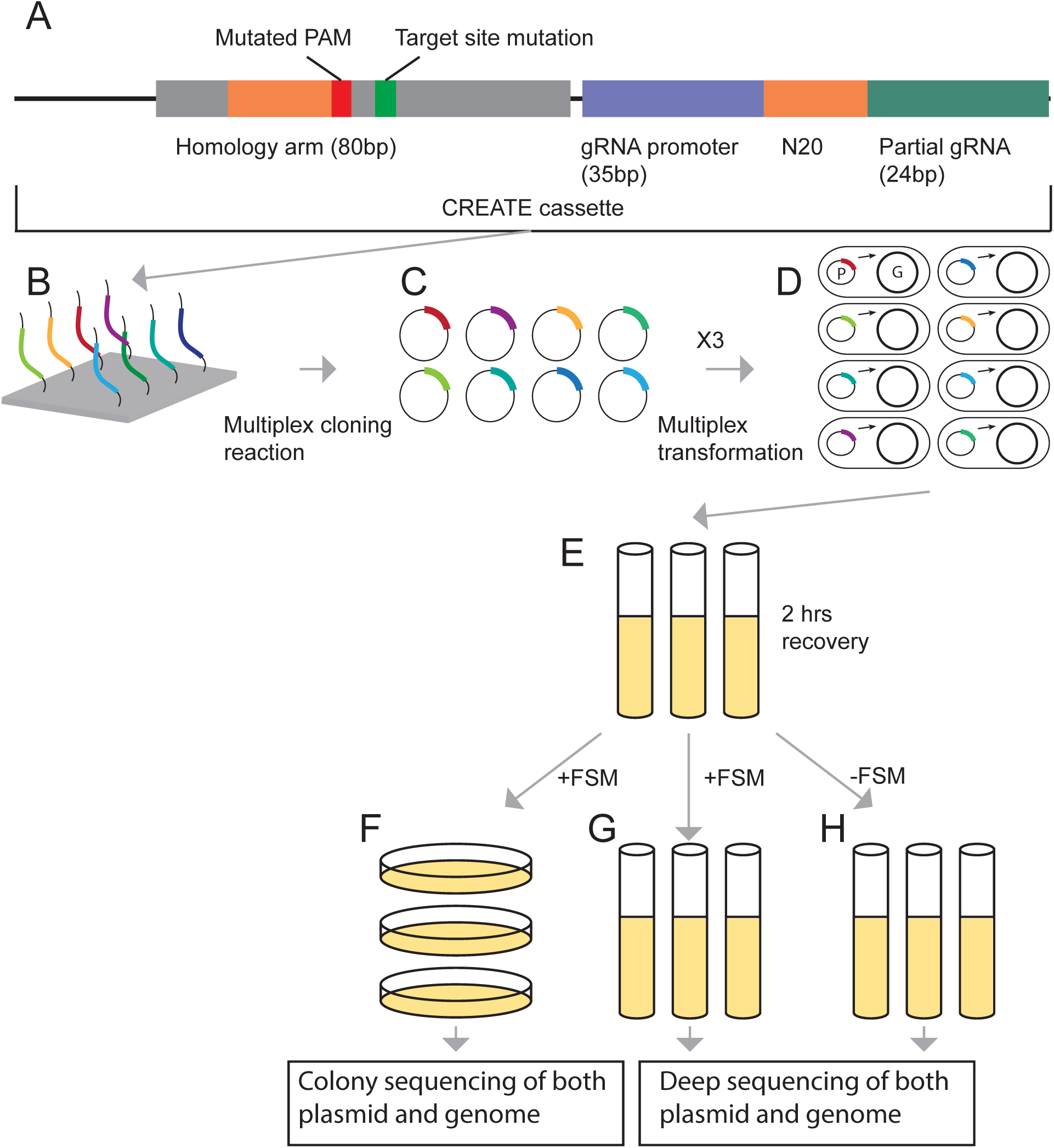
A schematic representation of the experimental procedure. A, an individual CREATE cassette includes a homology arm, harboring both the target site and adjacent mutations, and the corresponding gRNA. B, 660 such cassettes were synthesized on an array and were inserted into a plasmid backbone in a single multiplex cloning reaction (C). D, Plasmid library was transformed in triplicate into E. coli cells with the transcription induction of Cas9 and the recombineering machinery. Following 2 hours of recovery (E), cells were either plated on FSM-containing plates (F), or cultured overnight with (G) or without (H) the presence of FSM. Samples for deep sequencing were taken from E, G, and H. For a more detailed CREATE cassette, please refer to Fig. S1.

## Results

The *ispC* target sites included amino acids within the NADPH, ligand and FSM binding sites as described previously (29, 30), and resulted in a total of 32 target amino acids (Materials and Methods). An additional site was randomly added for control purposes, adding to a total of 33 sites (Fig. 2A). Each site was completely saturated to all remaining 19 amino acids and additionally to the native amino acid for control, resulting in a 660-mutant library. Each CREATE cassette harbors a mutated target site, an adjacent silently mutated PAM and a corresponding gRNA as depicted in Figure 1A and S1. The 660 cassette library was synthesized on an Agilent Array, together with other CREATE oligos designed for different experiments, as previously described (14). Genome-edited library cells were prepared in triplicates and allowed to recover for two hours. Library cells were incubated overnight either with or without 100uM FSM as well as plated on agar plates containing FSM (100uM).

**Fig. 2.**
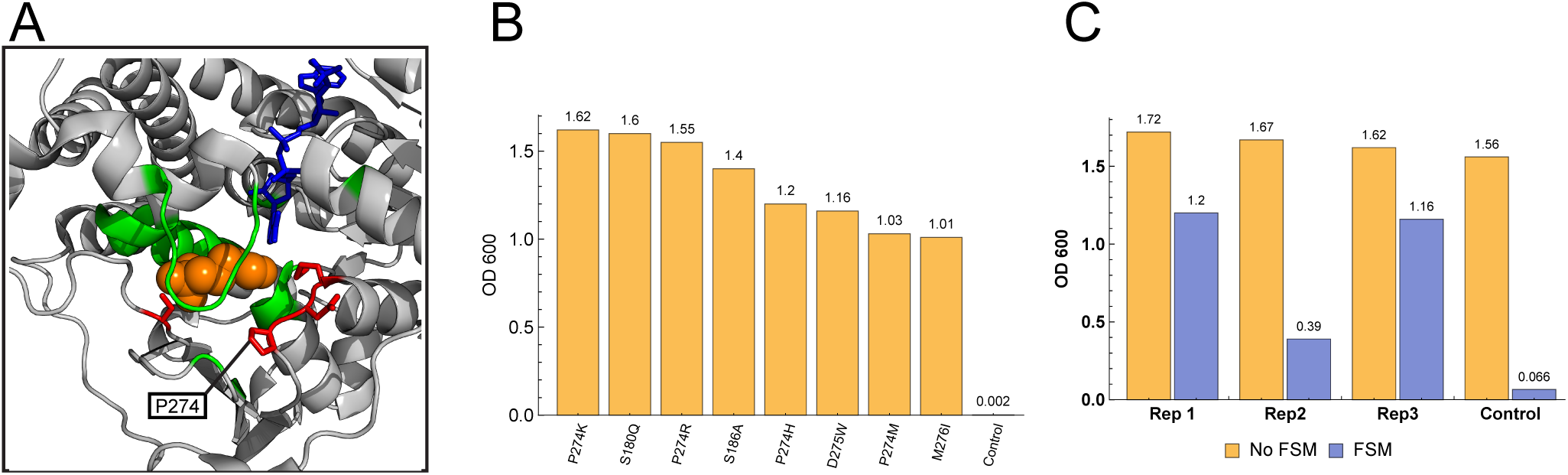
A, the DXR target sites. The DXR structure is shown in gray (PDB #1q0l). Red residues represent mutated sites enriched following FSM incubation, while the green amino acids represent the rest of the library. FSM is colored orange, and NADPH is in blue. B, eight different mutations were isolated from FSM-resistant colonies (step F in Fig.1). The CREATE plasmids were isolated, retransformed into fresh E. coli cells and treated with FSM. Optical density was measured following 16 hours of incubation. C. library cells growth in the presence of FSM. Optical density was measured 16 hours following FSM addition. Results correspond to steps G and H in Fig. 1. Control cells were edited in the E. coli galK gene which is not related to growth in LB or FSM resistance.

## FSM-resistant colonies

24 colonies were recovered from the FSM selection plates and were subjected to both CREATE plasmid and *IspC* genomic sequencing. Sequencing results are shown in Figure S2. Eight unique mutations in four sites were identified and confirmed genomically (Fig. 2A). The genomically edited colonies included CREATE plasmids that span between 100% identity to the mutational design, to four single base mismatches, indicating that CREATE editing can be robust to a small number of mutations within the homology arm. These errors might arise in the synthesis, amplification or cloning steps.

Out of the nine colonies (37.5%), that were not edited, plasmid sequencing revealed that seven had crossovers between the homology arm and the gRNA, and two had very large deletions (Fig. S2). Crossovers might occur during multiplex PCR library amplification (31) and result in a chimeric cassette. A mismatch between the homology arm and the gRNA portions of a CREATE cassette will likely result in cell death since there is no repairing template. Indeed, all of the crossovered gRNAs displayed additional gRNA spacer mutations, most probably rendering the gRNA ineffective (Fig. S2). These wild-type resistant colonies may have adapted to resist FSM via different mechanisms (see Discussion).

Cells harboring each of the eight individual mutations were cultured overnight, and the CREATE plasmid was extracted and re-transformed to new electrocompetent cells. Each culture, now harboring a single point mutation was subjected to 100uM FSM, similarly to the original selection. All mutations grew to higher optical density than control cells, validating that these mutations indeed confer FSM resistance. (Fig. 2B)

## Culture selection

The library cells in the selected liquid cultures grew to a larger optical density than control cells that were edited in the *galK* gene, a non-essential and non-DXP pathway gene (Fig. 2C). It should be noted that incubation times over 16 hours resulted in growth at the control tube, implying that adaptive events ultimately occur, as previously reported (see Discussion).

## CREATE Plasmid deep sequencing

The CREATE plasmids were extracted from all cultures (Fig. 1 E, G, H), the editing cassettes were PCR amplified and amplicons were subjected to deep sequencing. Sequences were filtered to include 99% identity to the designs, to ensure high editing confidence and adequate mutant identification. Total usable reads summed to about 3.36x10^6^ reads. Samples prior to selection included 630 different CREATE cassettes, constituting 95.4% of the designed library.

All three repeats correlate with each other in a statistically significant manner (Fig. S3A-C). To determine whether the library cells grew in the presence of FSM thanks to a specific set of mutations, enrichment analysis of the selected vs. the non-selected overnight cultures was performed (Fig. 1 G, H). To establish which mutations were significantly enriched, the average enrichment of the silent mutations was calculated as a baseline (see Materials and Methods). Mutations in proline 274 were highly enriched in all three replicates, indicating it may be a significant player in FSM resistance, in line with the results from individual colonies (Fig. S4). Importantly, a P274K mutation was common to all three repeats (Fig. S4). All mutations in the control site, G14, were within the error boundaries.

## Genomic sequencing of enriched mutants

Targeting a single gene allows measuring the edited genomes directly, in addition to indirect measurements via the CREATE plasmid. Since the *IspC* gene is longer than the maximal read length of the MiSeq platform (IspC is 1197 bp in length while the maximal read length using an Illumina Miseq is currently 600 bases.), the gene was amplified from the liquid cultures, and we used the Nextera XT (Agilent) kit to fragment the gene to smaller segments, resulting in less sequencing depth (See Discussion). Overall, a total of about 452,000 usable reads were obtained. Still, all three replicates significantly correlated with each other, although to a lesser extent than the plasmid sequencing (Fig. S3, D-F). Sequence analysis showed enriched mutations in all repeats and confirmed the plasmid results with P274 being highly mutated (Fig. 3). This analysis led to many more statistically-significant enriched mutations, probably due to the differences in the mean and standard deviation calculations (see Materials and Methods). Still, the mutagenesis effects of the control site, G14, were within the error bounds (Fig. S5). Both analyses agree on the majority of mutations within the repeats: The intersections between the significant plasmid and genomic enrichments are P274K for repeat 1, P274K, L230W, P274H and P274M for repeat 2, and P274K and S186Q for repeat 3. Interestingly, while P274K is common to all repeats, both repeats 2 and 3 include unique mutations that are shared in both sequencing approaches (L230W and S186Q, respectively). These differences between repeats may be due to unequal input DNA during library preparation or due to stochastic dynamics of cells survival following electroporation and recovery and were the rationale behind the decision to analyze every repeat individually. In addition to the intersect mutations, D275W and M276E/T were shown significant enrichments as well. Apart from L230W, all sites were also found in the colonies sequencing (Fig. 2B). Plotting the genomic enrichment vs. the plasmid enrichment values for every repeat showed that both sequencing approaches agree regarding resistant mutants within repeats and also between repeats, to some extent (Fig. S6).

**Fig. 3.**
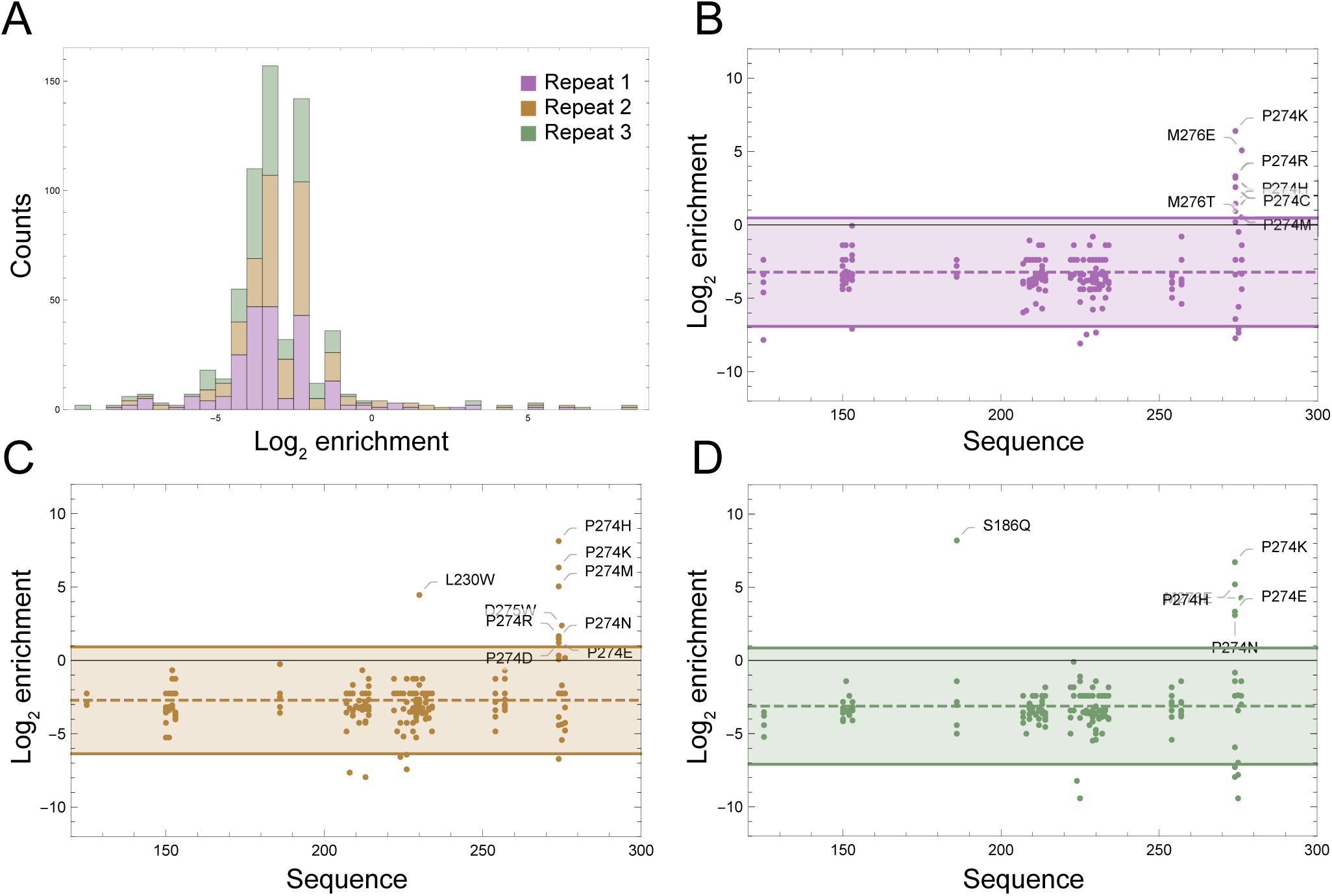
Enrichment analysis of library cells at the genomic level. A, enrichment histogram of all three repeats. B-D, Enriched mutants from repeats 1-3, respectively. The X axis represents the DXR sequence. The dashed line represents the average of all data points, and the solid lines are two standard deviations from the average. Mutants with enrichment values exceeding two standard deviations are labeled. For clarity, the randomly selected internal control (G14) was omitted and can be found in Fig. S5. A similar analysis was also performed using the CREATE reporter plasmid and can be found in Fig. S4.

## The P274 site

Mutations in P274 were found to be highly enriched in all FSM-treated repeats and the FSM-resistant colonies, suggesting that these mutations may confer FSM resistance while still retaining normal enzymatic activity. Figure 4 focuses on the genomic saturation data of this residue, organized by the average enrichment values of the three repeats. P274 in its three-dimensional context is shown in Figure S7. The identity of the enriched mutations indicates that a replacement of the proline with a charged amino acid increases the chances for resistance while replacing it with another hydrophobic residue does not generally result in enrichment. Interestingly, four out of the six most adaptive mutations require at least two base changes (highlighted in yellow), while the least adaptive mutations are enriched with single base transitions (red). This is in line with other observations, suggesting that highly adaptive mutants require a significant change in chemical properties, which are less accessible via single base change due to the conservative structure of the genetic code (14, 32–34) and are harder to isolate using error-based methods.

**Fig. 4.**
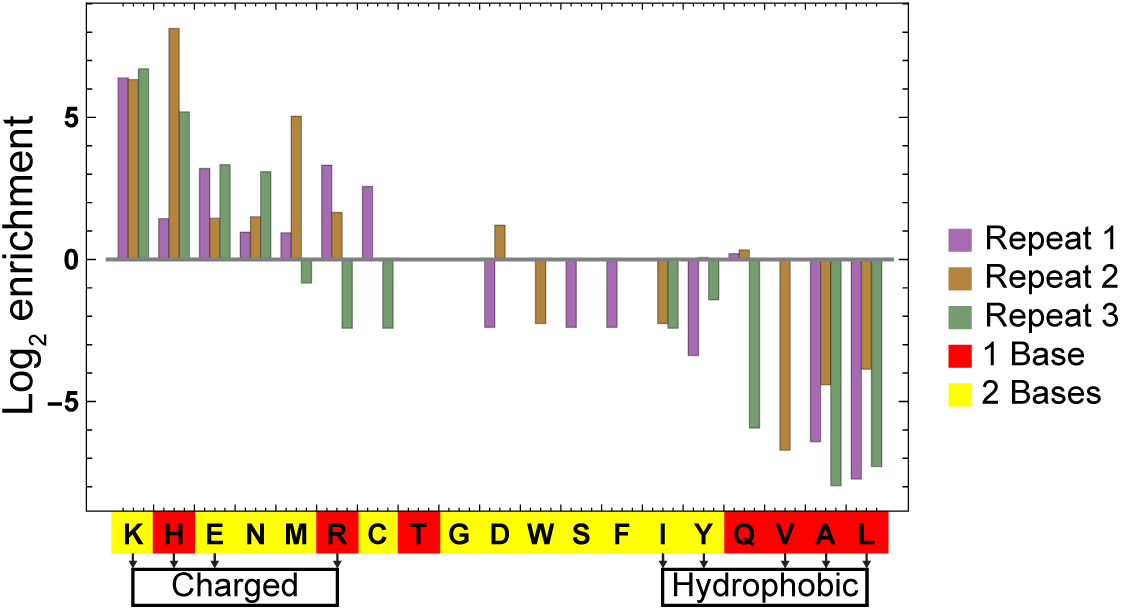
Saturation data of the P274 residue. The X axis represents the final amino acids replacing the wild-type proline and are ordered according to the average value of the three repeats. Y axis represents the enrichment values of each mutant. Red and yellow colors indicate whether the new amino acid requires one or two nucleotide changes in the wild-type codon, respectively. Note that the enriched (FSM resistant) mutants are enriched with charged amino acids, requiring two base changes, while the diluted mutants are a result of replacing a proline with another hydrophobic amino acid, needing a single base change only.

## Discussion

Drug resistance is an emerging concern for human health, affecting millions of patients worldwide. Drug resistance spans from aggressive multidrug-resistant bacteria in hospitals to acquired resistance of cancer cells, to resistance to infectious diseases such as malaria. Hence, the ability to predict the potential emergence of resistance mutations and their relative probability before they rise in the clinic is important for future drug design. Here, we applied the CREATE technology to generate a relatively small genomical saturation mutagenesis library of the *IspC* gene, with the aim to isolate mutations that confer resistance to its specific inhibitor, FSM.

The CREATE approach enables to edit target genomes rapidly, efficiently and systematically, leaving only a silent scar at an adjacent PAM site (14). Genome editing techniques are sometimes preferred over plasmid-based engineering since it is more biologically relevant, as expression from the plasmid alters the expression level and introduces bias. Another advantage of directly mutating the genome is the lack of a wild-type background allele that may potentially interfere with results interpretation.

Sequencing individual FSM-resistant colonies revealed that 100% match to the CREATE cassette design is not necessary, and up to four single base deletions (at least) are tolerated. Most unedited colonies harbored a chimeric cassette, with non-matching homology arms and gRNAs. While such errors are common in multiplex PCR reactions, there are several approaches to reduce them to a minimum (31, 35). Such crossovers, if not further mutated will result in cell death thanks to the gRNA-induced genome digestion with no available editing template and will be eliminated from the cell population. However, further mutations in the gRNA spacer region (as seen in all wild-type colonies, Figure S1) are a concern since these will not induce any double-strand breaks and might proliferate significantly faster than edited cells which require some recovery time prior to further growth. Hence, increasing the cassette design integrity is of importance. Here, we filtered the deep sequencing data to include cassettes that are 99% homologous to the original designs. This filtering approach allows up to only two point mutations within a cassette and removes all promiscuous cassettes from further analysis.

While CREATE was designed to provide indirect information on the enriched mutations via the editing cassette sequence at the plasmid level to enable genome-wide editing, targeting a single gene enables the its direct sequencing. Both direct and indirect sequencing were in general agreement and identified the most enriched mutants at the P274 site (Figures 3 and S4). Either method has advantages and disadvantages: First, the CREATE plasmid sequencing was much deeper, with significantly more relevant reads than the genomic sequencing. This is due to the sequencing method used for every run: while the CREATE cassette was designed to fit into a single deep sequencing read (300bp), the *E. coli IspC* gene is significantly longer, requiring a different sequencing approach. We used the Nextera XT DNA Library Preparation Kit (Illumina) that involves fragmenting of the PCR-amplified genomic segment. Since every gene copy was mutated in a single site, most fragments were wild-type and were filtered out at the analysis phase. In contrast, every read of the CREATE plasmid sequencing reaction contained relevant mutant information. However, CREATE plasmid reads might include false positive reads, correlating with the selection strength. Our colony sequencing resulted in some wild-type genomic *IspC*, indicating background adaptation. Indeed, unlike trimethoprim resistance that can be acquired mostly by mutations in its target, DHFR (36–38), FSM resistance may arise via other mechanisms. Expression of the efflux pump coded by the *E. coli* FSM-resistance gene, *fsr*, was shown to confer resistance (39), as well as deletions of the adenylate cyclase (*cya*) and the glycerol-3-phosphate transporter, *glpT* genes (40). Such adaptation events might have occurred, allowing cells to resist FSM, regardless of a mutation within the *IspC* gene. Hence, adapted cells, either harboring non-relevant mutations, or ones that failed to be edited by the CREATE mechanism and carry CREATE plasmids indicating for non-existent mutations are expected to contribute to noisy results. False positives may be identified and ignored by isolating mutations that are consistently enriched in several repeats, such as the case with the P274K mutation. Nevertheless, since both direct and indirect approaches agree on the most enriched mutants, CREATE is amenable for use in relatively weak selections such as FSM.

Mutations in proline 274 were highly enriched following FSM treatment. Adaptive mutations include charged residues which represent four out of the top six (K, H, E, and R) while most diluted mutations are composed of hydrophobic amino acids. Replacing proline with another hydrophobic amino acid may either not alter FSM binding at all, or disrupt the DXR structure to the point that it interferes with its essential enzymatic activity.

The P274 mutations that confer increased resistance to FSM mostly require more than a single base substitution within the codon, while the least adaptive mutations are mainly accessible via a single base replacement. This is in line with previous reports that in some cases the highest peaks in the mutational landscape are inaccessible when limited to single bases changes within a codon (14, 32–34). The reason for this is that sometimes dramatic phenotypic changes require a considerable difference in structural or chemical properties, which are buffered by the genetic code structure.

In line with the mention above, an analysis of the mutational fitness values as a function of the minimal number of required base change, showed a similar global trend (Fig. S8), emphasizing the advantage of saturation mutagenesis methods over random-based approaches that are unable to search the complete sequence space. It is important to note that we have used the minimum number of base change metric instead of a codon-specific number of substitutions, so the data is representative across organisms utilizing different codon usage. Having two or three base changes within a codon may occur either simultaneously (given enough genomes and high error rates), or sequentially (given enough time and depending on the fitness contribution of each individual step). Nevertheless, these are not expected to emerge in laboratory timescales and are less likely to occur naturally.

To date, a single mutation, S222T, was reported to confer FSM resistance in E. coli (28). While S222 was targeted in our library, it was not detected as enriched following FSM selection, but rather it was depleted between the two hours recovered sample and the overnight, unselected culture. This discrepancy may be explained by the differences between the two approaches: while Armstrong et al. isolated the mutant by colony picking, here, the selection was made in culture, at the population level, adding an element of competition between resistant clones during recovery as well as during overnight culture. Indeed, the *K_m_*of the S222T mutant was shown to be significantly increased, reducing the affinity of the enzyme to its substrate, DOXP (28). We have reconstructed the S222T mutation using the CREATE approach and validated its FSM-resistance properties.

The ability to perform rapid and systematic sequence to activity mapping in multiplex facilitates the isolation of drug-specific resistant mutants before they occur in the field. Armed with such knowledge, drug developers may derive drugs that it is less likely to develop mutational acquired resistance for. Moreover, such drug candidates may be tested using similar libraries increasing the efficiency of the development cycle. While this study was made on the *E. coli IspC* gene, the results presented here are of relevance for other, more clinically relevant pathogens. Multiple alignments of the complete list of reviewed DXR genes in Uniprot (41), spanning from bacteria through protozoa to plants, show high conservation values of the identified sites: P274 is conserved in 87.8% of the proteins while S186, D275, and M276 are 100%, 95.9% and 99.3% conserved, respectively. Importantly, all these sites are identical both in *E. coli* and *Plasmodium falciparum*, making these results especially relevant for antimalarial drug development.

## Materials and methods

### Plasmids and strains

plasmid coding for Cas9 was previously described (42). The CREATE backbone plasmid was described in the first CREATE paper (14). The recombineering plasmid used in this study was pSim5 (43). All experiments were performed using the standard K12 MG1655 strain.

### CREATE cassette design

a CREATE cassette is 200bp, harboring two elements: (1) an editing segment, or homology arm that correspond to a genomic region containing the target site. The target site is altered to translate to the amino acid of interest, and an adjacent PAM site is silently mutated to eliminate Cas9 recognition, but to preserve the amino acid sequence. (2) A corresponding gRNA, targeting the mutated PAM site. For a sequence example refer to Fig. S1. The complete library design can be found as a supplementary file for this article.

### Target sites

Target sites are based on previous structural analyses and represent portions of the NADPH, ligand, and FSM interacting amino acids (29, 30). The library includes the following sites: 125, 150, 151, 152, 153, 186, 207, 208, 209, 210, 211, 212, 213, 214, 222, 223, 224, 225, 226, 227, 228, 229, 230, 231, 232, 233, 234, 254, 257, 274, 275, 276. Amino acid number 14 was randomly added to the library as an internal control.

### Library preparation

For a detailed protocol refer to Garst et al. (14). *IspC* library oligos were purchased from Agilent and were received pooled with other CREATE libraries. DNA was purified using acrylamide gel electrophoresis. The *IspC* library was amplified using its unique primers, was cloned into a plasmid backbone and was transformed into electrocompetent cells. Library plasmids were extracted (miniprep kit, Qiagen) and were transformed into electrocompetent, heat-shocked target K12 MG1655 cells that were already harboring both pSim5 and Cas9 plasmids. For control, a *galK* (a Y145 to a stop codon mutation) CREATE plasmid was also transformed to separate cells. The galK edited cells were also used to assess editing efficiency using a pink/white colorimetric assay (44, 45) Cells were recovered in LB with carbenicillin (CREATE plasmid), kanamycin (Cas9), and arabinose (Cas9 induction) for two hours.

### Selection

All selection experiments were performed with 100uM FSM (Molecular Probes, catalog number: F23103). Recovered cells were divided into three groups and were either plated on FSM-containing plates, or cultured in fresh media with or without FSM, and with carbenicillin, kanamycin, and arabinose (Fig. 1). Cells were incubated overnight at 37°C following optical density measurements and colony picking.

### Deep sequencing

For CREATE plasmid sequencing, the plasmid was extracted from the cultures (steps E, G, and H in Fig. 1), and the editing cassettes were PCR-amplified, with each sample harboring a different sequence barcode, as shown before (14). Amplicons were purified and pooled. For the genomic sequencing, Cells were boiled for 5 minutes and the complete IspC gene was amplified from the different samples, including 200 base pairs up- and downstream from the gene using the following primers: (forward - GCACTGTTGAAAGATAAAGAGATCAGCG and reverse - CTGGCAATTTTTCGCTAAGTGGTTGAG). Samples were digested and barcoded using the Nextera XT kit (Illumina). Both samples were sequenced using a MiSeq deep sequencer (Illumina) by the University of Colorado Boulder sequencing core.

### Plasmid-based CREATE tracking analysis

Processing of high-throughput sequencing reads and query matching: High-throughput sequencing of CREATE plasmids were performed using an Illumina MiSeq 2x150 paired-end reads run. Reads were demultiplexed according to an experiment-specific unique barcode, allowing a maximum of 3 mismatches, and then merged using the PANDAseq assembler (v2.10). Merged reads were matched to the database of all designed cassettes using the usearch_global algorithm (v9.2.64), with an identity threshold of 90% and minimal alignment length of 150 bp (75% of total). 40 hits were allowed for each query, which were subsequently sorted by percent identity and the best-matching cassette was chosen. To generate read counts for each designed cassette, only reads that had a full alignment and an identity higher than 99% were used.

Data filtering and enrichment analysis: Data frames of read counts were generated and processed using the Pandas data analysis python package (v0.20.2). Final read counts for each cassette after FSM selection were compared against counts for a 2 hours growth baseline and overnight growth baseline (without selection). Enrichment scores were calculated as the logarithm (based 2) of the ratio of frequencies between post-selection (FSM) to pre-selection for each individual replicate. Frequencies were determined by dividing the read counts for each variant to the total experimental reads. Since low-count variants are subject to counting error, variants with initial counts that sum up to less than 100 across all three replicates were not included in the analysis. The enrichment scores were used to rank the fitness contributions of all variants under FSM selection in each individual replicate. To assess significance, the average of enrichment scores for all synonymous mutations included in the library were considered (i.e. average µ of wild-type enrichment). Bootstrap analysis (resampled with replacement 20,000 times) were performed to obtain a 95% confidence interval for the wild-type enrichment average µ). Variants were considered as significantly enriched under FSM selection if their enrichment scores were at least µ ± 2* (i.e. p-value 0.05 assuming a normal distribution of enrichment scores).

### Genomic ispC variant sequencing analysis

Data processing and variant calling: Following standard quality filtering and demultiplexing the reads, variants were called using a k-mer strategy. Briefly, a database was generated with all designed ispC mutations. Then, a unique 10-mer sequence was searched in the database for each variant using custom python scripts. It should be noted that 23 cassettes, all encoding synonymous mutations, did not contain any unique k-mers and were not included in the analysis. After the list of unique k-mers were generated, variants were called and counted using the grep command for Unix operating systems. Enrichment analysis: Variant counts post-selection (FSM) were compared to pre-selection counts (2 hours or overnight) using the Pandas data analysis python package (v0.20.2). Similarly to the analysis performed for CREATE plasmids, enrichment scores were calculated as the logarithm (based 2) of the ratio of frequencies between post-selection (FSM) to pre-selection for each individual replicate, with frequencies determined by dividing the read counts for each variant to the total experimental reads. To assess significance, the mean values of a complete comparison was calculated and significance was set to 2 standard deviations from the calculated mean.

### Mutants validation

Single colonies, successfully growing on FSM were picked and cultured overnight. CREATE plasmids were extracted and retransformed to fresh, heat-shocked, electrocompetent K12 MG1655 cells, already harboring the Cas9 and Psim5 plasmids. Following the same procedure described in *Library preparation*, growth in the presence of FSM was assayed by measuring optical density after overnight growth.

## Acknowledgments

We would like to thank Yael David for her helpful insights. This work was funded both by the US Department of Energy Grant No. DE-SC008812 and by the National Institute of Allergy and Infectious Diseases (NIAID) at the NIH, grant number 1R21AI128296-01A1

## Author contributions

G.P., A.D.G, A.C. and R.T.G. designed research; G.P., E.J.O. and A.C. performed research; G.P., M.C.B. and A.C. analyzed data; and G.P and R.T.G. wrote the paper.

